# The Structural Basis of Rubisco Phase Separation in the Pyrenoid

**DOI:** 10.1101/2020.08.16.252809

**Authors:** Shan He, Hui-Ting Chou, Doreen Matthies, Tobias Wunder, Moritz T. Meyer, Nicky Atkinson, Antonio Martinez-Sanchez, Philip D. Jeffrey, Sarah A. Port, Weronika Patena, Guanhua He, Vivian K. Chen, Frederick M. Hughson, Alistair J. McCormick, Oliver Mueller-Cajar, Benjamin D. Engel, Zhiheng Yu, Martin C. Jonikas

## Abstract

Approximately one-third of global CO_2_ fixation occurs in a phase separated algal organelle called the pyrenoid. Existing data suggest that the pyrenoid forms by the phase-separation of the CO_2_-fixing enzyme Rubisco with a linker protein; however, the molecular interactions underlying this phase-separation remain unknown. Here we present the structural basis of the interactions between Rubisco and its intrinsically disordered linker protein EPYC1 (Essential Pyrenoid Component 1) in the model alga *Chlamydomonas reinhardtii*. We find that EPYC1 consists of five evenly-spaced Rubisco-binding regions that share sequence similarity. Single-particle cryo-electron microscopy of one of these regions in complex with Rubisco indicates that each Rubisco holoenzyme has eight binding sites for EPYC1, one on each Rubisco small subunit. Interface mutations disrupt binding, phase separation, and pyrenoid formation. Cryo-electron tomography supports a model where EPYC1 and Rubisco form a co-dependent multivalent network of specific low-affinity bonds, giving the matrix liquid-like properties. Our results advance the structural and functional understanding of the phase separation underlying the pyrenoid, an organelle that plays a fundamental role in the global carbon cycle.

## Main Text

The CO_2_-fixing enzyme Rubisco drives the global carbon cycle, mediating the assimilation of approximately 100 gigatons of carbon per year^1^. The gradual decrease of atmospheric CO_2_ over billions of years^2^ has made Rubisco’s job increasingly difficult, to the point where CO_2_ assimilation limits the growth rate of many photosynthetic organisms^3^. This selective pressure is thought to have led to the evolution of CO_2_ concentrating mechanisms, which feed concentrated CO_2_ to Rubisco to enhance growth^4^. Of these mechanisms, the most poorly understood relies on the pyrenoid, a phase separated organelle^5^ found in the chloroplast of nearly all eukaryotic algae and some land plants (Fig. 1a, b)^6,7^. The pyrenoid enhances the activity of Rubisco by clustering it around modified thylakoid membranes that supply Rubisco with concentrated CO_2_^8,9^.

**Fig. 1.**
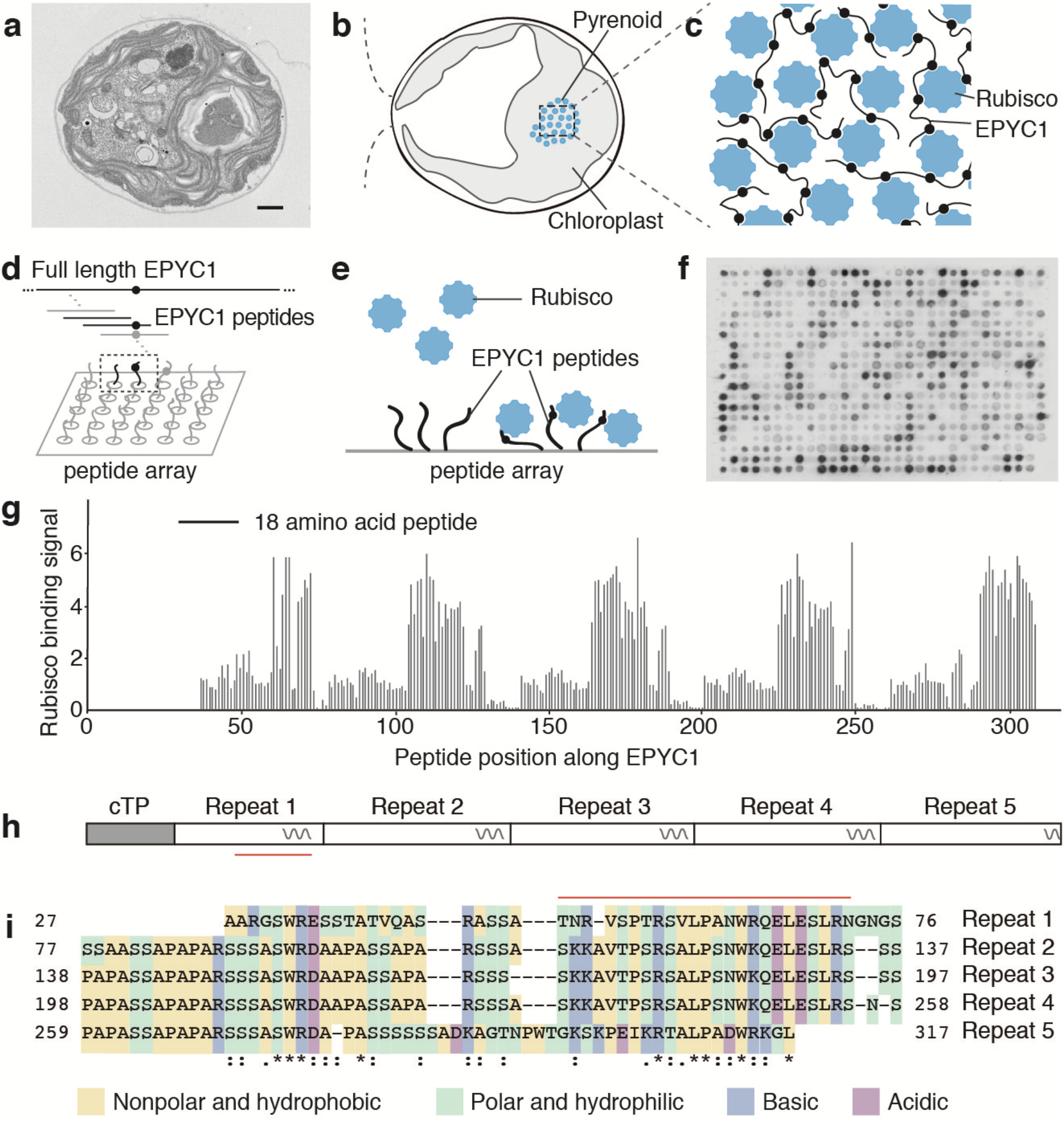
EPYC1 consists of five tandem sequence repeats, each of which contains a Rubisco-binding region. **a**, Transmission electron microscopy (TEM) image of a Chlamydomonas cell. Scale bar = 1 μm. **b**, Cartoon depicting the chloroplast and pyrenoid in the image shown in panel a. The blue dots indicate the location of Rubisco enzymes clustered in the pyrenoid matrix. **c**, We hypothesized that pyrenoid matrix formation is mediated by multivalent interactions between Rubisco and the intrinsically disordered protein EPYC1. **d**, We designed an array of 18 amino acid peptides tiling across the full length EPYC1 sequence. **e**, Incubation of the array with purified Rubisco allows identification of peptides that bind to Rubisco. **f**, Image of the Rubisco binding signal from the peptide tiling array. **g**, The Rubisco binding signal was quantified and plotted for each peptide as a function of the position of the middle of the peptide along the EPYC1 sequence. The initial 26 amino acids of EPYC1 correspond to a chloroplast targeting peptide (cTP), which is not present in the mature protein^12^. Results are representative of three independent experiments. **h**, The positions of EPYC1’s five sequence repeats are shown to scale with panel g. Predicted α-helical regions are shown as wavy lines. **i**, Primary sequence of EPYC1, with the five sequence repeats aligned. In panels h and i, the region used for structural studies (EPYC149-72) is indicated by a red line.

For decades, the mechanism for packaging the Rubisco holoenzyme into the pyrenoid remained unknown. Recent work showed that in the leading model alga *Chlamydomonas reinhardtii* (Chlamydomonas hereafter), the clustering of Rubisco into the pyrenoid matrix requires the Rubisco-binding protein EPYC1^10^. EPYC1 and Rubisco are the most abundant components of the pyrenoid and bind to each other. Moreover, combining purified EPYC1 and Rubisco together produces phase-separated condensates^11^ that mix internally at a rate similar to that observed for the matrix *in vivo*^5^, suggesting that these two proteins are sufficient to form the structure of the pyrenoid matrix. The sequence repeats within EPYC1 and eight-fold symmetry of the Rubisco holoenzyme led us to hypothesize that EPYC1 and Rubisco each have multiple binding sites for the other, allowing the two proteins to form a co-dependent condensate (Fig. 1c)^10^.

Here, we determined the structural basis that underlies the EPYC1-Rubisco condensate. Using biophysical approaches, we found that EPYC1 has five evenly spaced Rubisco-binding regions that share sequence homology and can bind to Rubisco as short peptides. We obtained a cryo-electron microscopy structure, which shows that each of EPYC1’s Rubisco-binding regions forms an α-helix that binds one of Rubisco’s eight small subunits via salt bridges and hydrophobic interactions. Mapping of these binding sites onto Rubisco holoenzymes within the native pyrenoid matrix indicates that the linker sequences between Rubisco-binding regions on EPYC1 are sufficiently long to connect together adjacent Rubisco holoenzymes. These discoveries advance the understanding of the pyrenoid, and provide a high resolution structural view of a phase-separated organelle.

## Results

### EPYC1 has five nearly-identical Rubisco-binding regions

We could not directly determine the structure of full-length EPYC1 bound to Rubisco because mixing the two proteins together produces phase separated condensates^11^. We thus aimed to first identify Rubisco-binding regions on EPYC1, and subsequently to use a structural approach to determine how these regions bind to Rubisco.

The intrinsically disordered nature of purified EPYC1^11^ led us to hypothesize that the Rubisco-binding regions of EPYC1 were short and could bind to Rubisco as peptides without a need for tertiary folds. Therefore, to identify EPYC1 regions that bind to Rubisco, we synthesized a peptide array consisting of 18 amino acid peptides tiling across the full length EPYC1 sequence (Fig. 1d), and probed this array with native Rubisco purified from Chlamydomonas cells (Fig. 1e, f).

Our tiling array revealed five evenly-spaced Rubisco-binding regions on EPYC1, each consisting of a predicted α-helix and an upstream region (Fig. 1g, h). We confirmed the binding regions using surface plasmon resonance (SPR; Extended Data Fig. 1b, c). Sequence alignment guided by the five binding regions revealed that mature EPYC1 consists entirely of five sequence repeats (Fig. 1i), in contrast to the previously defined four repeats and two termini^10^ (Extended Data Fig. 1a). Our alignment indicates that the previously defined EPYC1 N- and C-termini, which at the time were not considered part of the repeats, actually share sequence homology with the central repeats.

The presence of a Rubisco-binding region on each of the previously defined EPYC1 repeats (Extended Data Fig. 1a) explains our yeast two-hybrid observations^12^ that a single EPYC1 repeat can interact with Rubisco, that knocking out the α-helix in an EPYC1 repeat disrupts this interaction, and that decreasing the number of EPYC1 repeats leads to a proportional decrease in EPYC1 interaction with Rubisco. It also explains our observations that decreasing the number of EPYC1 repeats leads to a proportional decrease in the tendency of EPYC1 and Rubisco to phase separate together^11^.

### EPYC1 binds to Rubisco small subunits

The sequence homology of the five Rubisco-binding regions suggests that each region binds to Rubisco in a similar manner. To identify the binding site of EPYC1 on Rubisco, we performed single-particle cryo-electron microscopy on a complex of Rubisco and a peptide corresponding to the first Rubisco-binding region of EPYC1 (Fig. 2a). We selected this region of EPYC1 because in preliminary experiments it had the highest affinity to Rubisco, which was still low by protein interaction standards (K_D_ ~3 mM; Extended Data Fig. 1d, e). This low affinity meant that millimolar concentrations of peptide were required to approach full occupancy of peptide bound to Rubisco, leading to challenges with peptide insolubility and high background signal in the electron micrographs. Despite these challenges, we successfully obtained a 2.62 Å structure of the complex (~2.9 Å EPYC1 peptide local resolution; Fig. 2, Extended Data Fig. 2 and 3; Extended Data Table 1). For reference purposes, we also obtained a 2.68 Å cryo-electron density map of Rubisco in the absence of EPYC1 peptide (Extended Data Fig. 2), which was nearly identical to the previously published X-ray crystallography structure^13^, with minor differences likely due to the absence of the substrate analog 2-CABP in the active site of Rubisco in our sample^14^ (Extended Data Fig. 4).

**Fig. 2.**
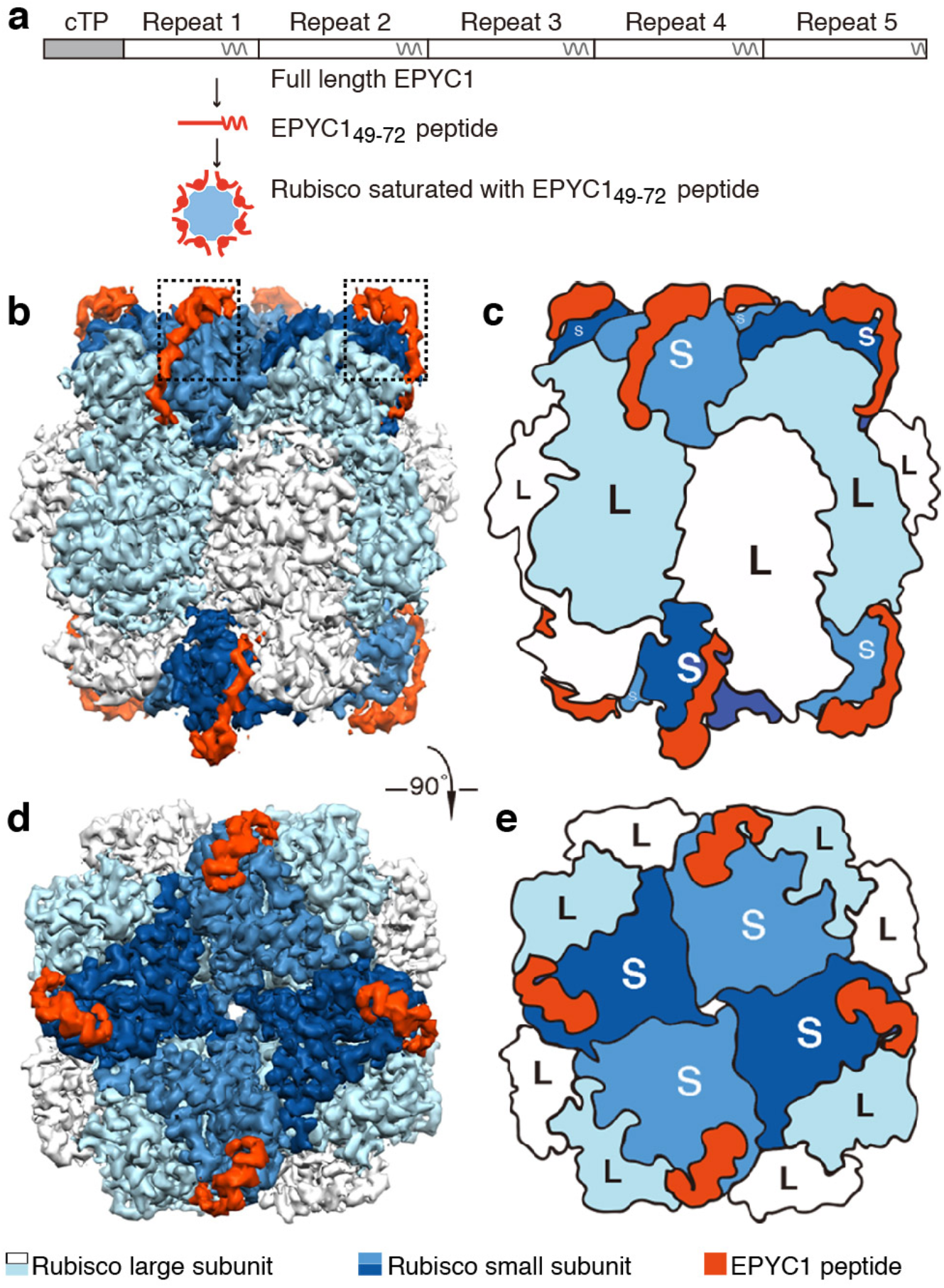
EPYC1 binds to Rubisco small subunits. **a**, Peptide EPYC149-72, corresponding to the first Rubisco-binding region of EPYC1, was incubated at saturating concentrations with Rubisco prior to single particle cryo-electron microscopy. **b-e**, Density maps (b, d) and cartoons (c, e) illustrate the side views (b, c) and top views (d, e) of the density map of the EPYC1 peptide-Rubisco complex. Dashes in panel b indicate regions shown in Fig.3a-3f.

**Fig. 3.**
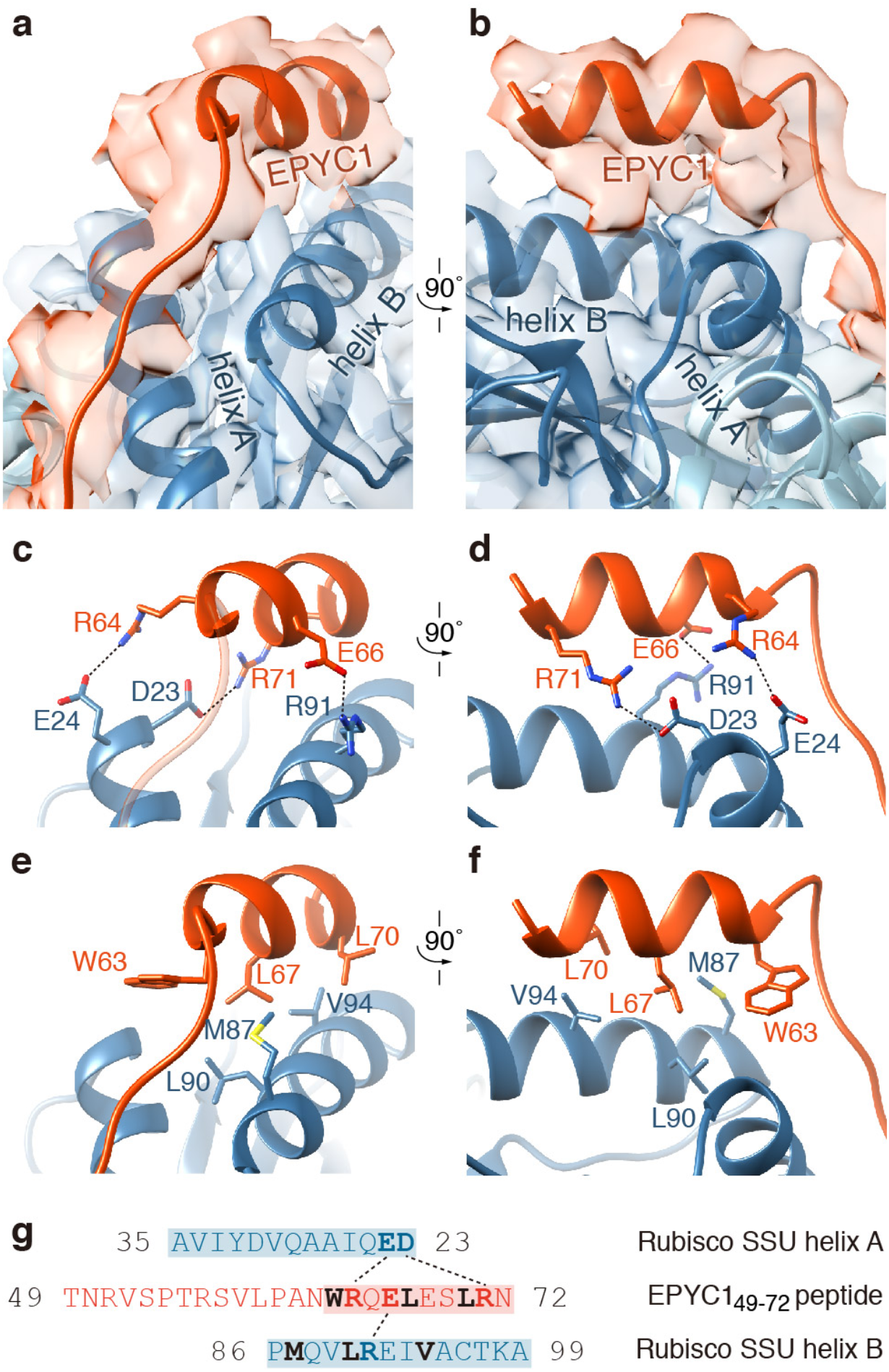
EPYC1 binds to Rubisco small subunit α-helices via salt bridges and a hydrophobic pocket. **a-b**, Front (a) and side (b) views of the EPYC1 peptide (red) bound to the two α-helices of the Rubisco small subunit (blue). **c-d**, Three pairs of residues form salt bridges between the helix of the EPYC1 peptide and the helices on the Rubisco small subunit. Shown are front (c) and side (d) views as in panel a and panel b. The distances from EPYC1 R64, R71 and E66 to Rubisco small subunit E24, D23 and R91 are 3.06 Å, 3.23 Å, and 3.13 Å, respectively. **e-f**, A hydrophobic pocket is formed by three residues of the EPYC1 peptide and three residues of helix B of the Rubisco small subunit. Shown are front (e) and side (f) views as in panel a and panel b. g, Summary of the interactions observed between the EPYC1 peptide and the two α-helices of the Rubisco small subunit. Helices are highlighted; the residues mediating interactions are bold; salt bridges are shown as dotted lines; residues contributing to the hydrophobic pocket are shown in black.

The Rubisco holoenzyme consists of a core of eight catalytic large subunits in complex with eight small subunits, four of which cap each end of the holoenzyme (Fig. 2b-e). In our structure, an EPYC1 peptide was clearly visible bound to each Rubisco small subunit, suggesting that each Rubisco holoenzyme can bind up to eight EPYC1s (Fig. 2b-e).

### Binding is mediated by salt bridges and a hydrophobic interface

The EPYC1 peptide forms an extended chain that sits on top of the Rubisco small subunit’s two α-helices (Fig. 3a, b). The structure explains our previous observations that mutations in the Rubisco small subunit α-helices disrupted yeast two-hybrid interactions between EPYC1 and the Rubisco small subunit^12^ and prevented Rubisco’s assembly into a pyrenoid *in vivo*^15^. The C-terminal region of the EPYC1 peptide (NWRQELESLRN) is well-resolved and forms an α-helix that runs parallel to helix B of the Rubisco small subunit (Fig. 3a, b). The peptide’s N-terminus extends the trajectory of the helix and follows the surface of the Rubisco small subunit (Fig. 2b-e, 3a-b and Extended Data Fig. 5a). The side chains of the peptide’s N-terminus could not be well resolved, suggesting that this region is more conformationally flexible.

Our atomic model based on the density map suggests that binding is mediated by salt bridges and a hydrophobic interface. Three residue pairs likely form salt bridges (Fig. 3c, d and g): EPYC1 residues R64 and R71 interact with E24 and D23, respectively, of Rubisco small subunit α-helix A, and EPYC1 residue E66 interacts with R91 of Rubisco small subunit α-helix B. Furthermore, a hydrophobic interface is formed by W63, L67 and L70 of EPYC1 and M87, L90 and V94 of Rubisco small subunit helix B (Fig. 3e-g).

### Interface residues are required for binding and phase separation *in vitro*

To determine the importance of individual EPYC1 residues for binding, we investigated the impact on Rubisco binding of every possible single amino acid substitution for EPYC1’s first Rubisco-binding region by using a peptide array (Fig. 4a) and SPR (Extended Data Fig. 5b). Consistent with our structural model, the peptide array indicated that EPYC1 salt bridge-forming residues R64, R71 and E66 and the hydrophobic interface residues W63, L67 and L70 were all required for normal EPYC1 binding to Rubisco. The strong agreement of our mutational analysis suggests that our structural model correctly represents EPYC1’s Rubisco-binding interface.

**Fig. 4.**
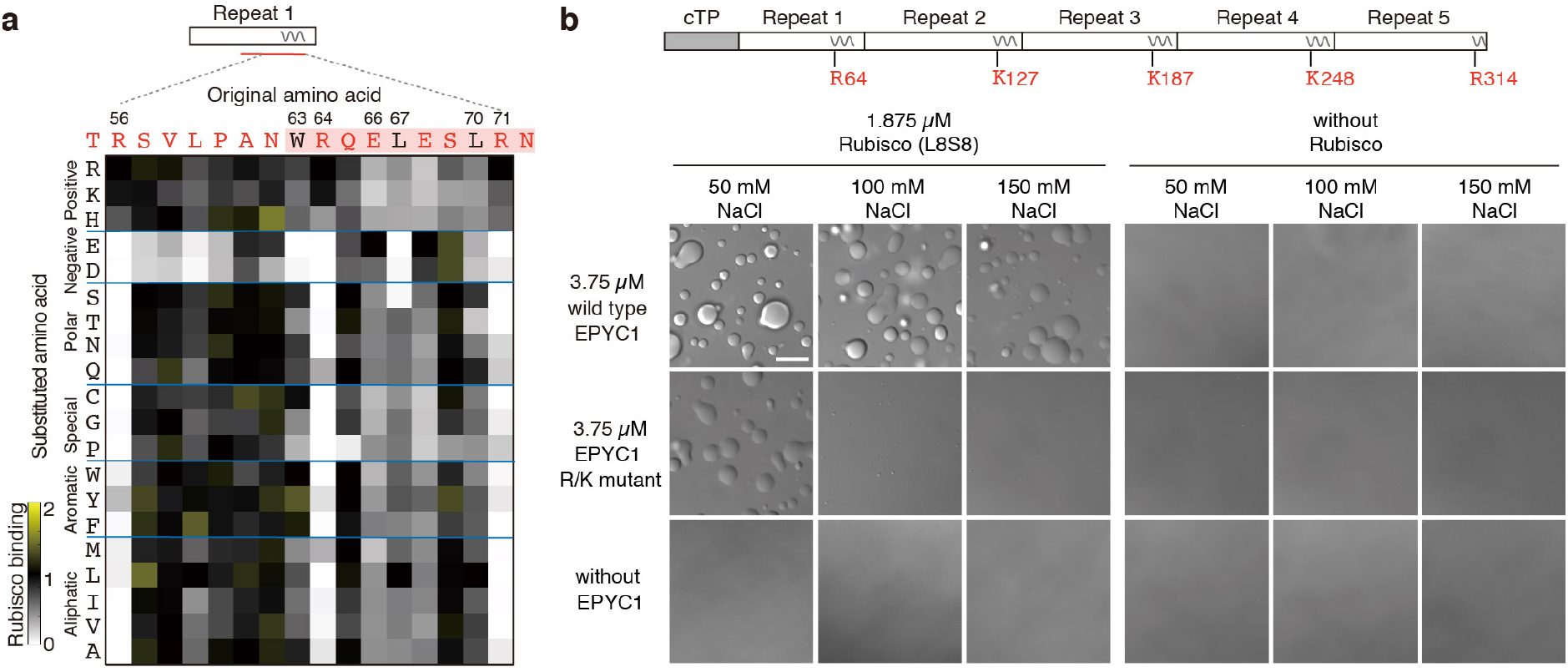
Interface residues on EPYC1 are required for binding and phase separation of EPYC1 and Rubisco *in vitro*. **a**, Rubisco binding to a peptide array representing every possible single amino acid substitution for amino acids 56-71 of EPYC1. The binding signal was normalized by the binding signal of the original sequence. **b**, The effect of mutating the central R or K in each of EPYC1’s Rubisco-binding regions on *in vitro* phase separation of EPYC1 with Rubisco. Scale bar = 10 μm.

To determine the importance of EPYC1’s Rubisco-binding regions for pyrenoid matrix formation, we assayed the impact of mutations in these regions on formation of phase separated droplets by EPYC1 and Rubisco *in vitro*. The phase boundary was shifted by mutating R64 in the first Rubisco-binding region and the corresponding K or R in the other four Rubisco-binding regions of EPYC1 (Fig. 4b and Extended Data Fig. 5c-e), suggesting that the Rubisco-binding regions mediate condensate formation.

### The binding interface is required for pyrenoid matrix formation *in vivo*

We validated the importance of Rubisco residues for binding to EPYC1 by yeast two-hybrid assays (Fig. 5a and Extended Data Fig. 6). Rubisco small subunit D23A mutation, which eliminates the charge of that residue, had a severe impact on Rubisco small subunit interaction with EPYC1, as expected from the contribution of that residue to a salt bridge with R71 of EPYC1. Likewise, E24A and R91A each showed a moderate defect, consistent with the contributions of those residues to salt bridges with R64 and E66 of EPYC1, respectively. Additionally, M87D and V94D, which convert hydrophobic residues to bulky charged residues, each had a severe impact on interaction, as expected from the participation of those residues in the hydrophobic interface. Combinations of these mutations abolished the interactions completely (Extended Data Fig. 6).

**Fig. 5.**
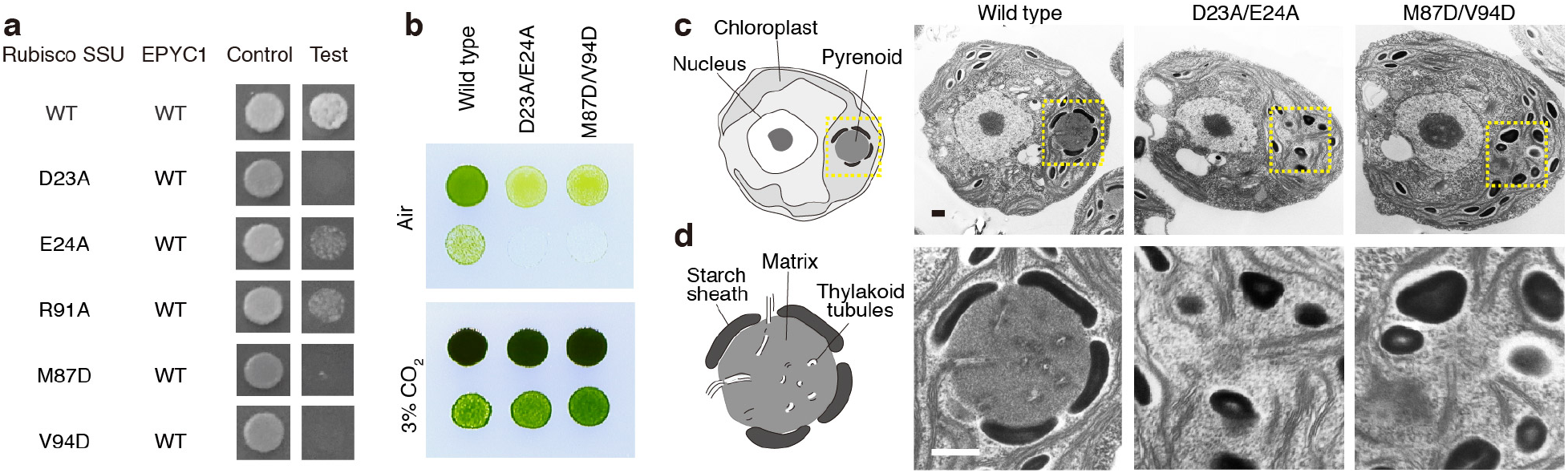
Interface residues on Rubisco are required for Yeast Two-Hybrid interactions between EPYC1 and Rubisco, and for pyrenoid matrix formation *in vivo*. **a**, The importance of Rubisco small subunit residues for interaction with EPYC1 was tested by mutagenesis in a yeast two-hybrid experiment. **b**, The Rubisco small subunit-less mutant T60 (*Δrbcs*) was transformed with wild-type, D23A/E24A or M87D/V94D Rubisco small subunits. Serial 1:10 dilutions of cell cultures were spotted on TP minimal medium and grown in air or 3% CO_2_. **c-d**, Representative electron micrographs of the whole cells (c) and corresponding pyrenoids (d) of the strains expressing wild-type, D23A/E24A, and M87D/V94D Rubisco small subunit. Dashes in panel c indicate regions shown in panel F. Scale bars = 500 nm.

To evaluate the importance of the binding interface *in vivo*, we generated Chlamydomonas strains with point mutations in the binding interface. Rubisco small subunit mutations D23A/E24A or M87D/V94D caused a growth defect under conditions requiring a functional pyrenoid (Fig. 5b, Extended Data Fig. 7a-b). Furthermore, the mutants lacked a visible pyrenoid matrix (Fig. 5c, d and Extended Data Fig. 7c), indicating that those Rubisco small subunit residues are required for matrix formation *in vivo*. The Rubisco mutants retained pyrenoid tubules^16^, as previously observed in other matrix-deficient mutants^10,17–19^.

Together, our data demonstrate that EPYC1’s Rubisco-binding regions bind to the Rubisco small subunit α-helices via salt-bridge interactions and a hydrophobic interface, enabling the condensation of Rubisco into the phase separated matrix.

### The spacing between EPYC1’s Rubisco-binding regions allows linking of adjacent Rubisco holoenzymes in the native pyrenoid matrix

The presence of multiple Rubisco-binding regions along the EPYC1 sequence supports a model where consecutive Rubisco-binding regions on the same EPYC1 polypeptide can bind to different Rubisco holoenzymes and thus hold them together to form the pyrenoid matrix. If this model is correct, we would expect that the ~40 amino acid “linker” regions between consecutive Rubisco-binding regions on EPYC1 (Fig. 1g, i) would be sufficient to span the distance between EPYC1-binding sites on neighboring Rubisco holoenzymes in the pyrenoid matrix. To test this aspect of the model, we combined our atomic structure of the EPYC1-Rubisco interaction with the precise positions and orientations of Rubisco holoenzymes within the pyrenoid matrix of native cells that we had previously obtained by *in-situ* cryo-electron tomography^5^ (Fig. 6a, b). We mapped the positions of EPYC1 binding sites onto Rubisco holoenzymes in the matrix and measured the distances between nearest neighbor EPYC1 binding sites on adjacent holoenzymes (Fig. 6c). The observed distances ranged from ~2 nm to ~7 nm, with a median distance of ~4 nm (Fig. 6d).

**Fig. 6.**
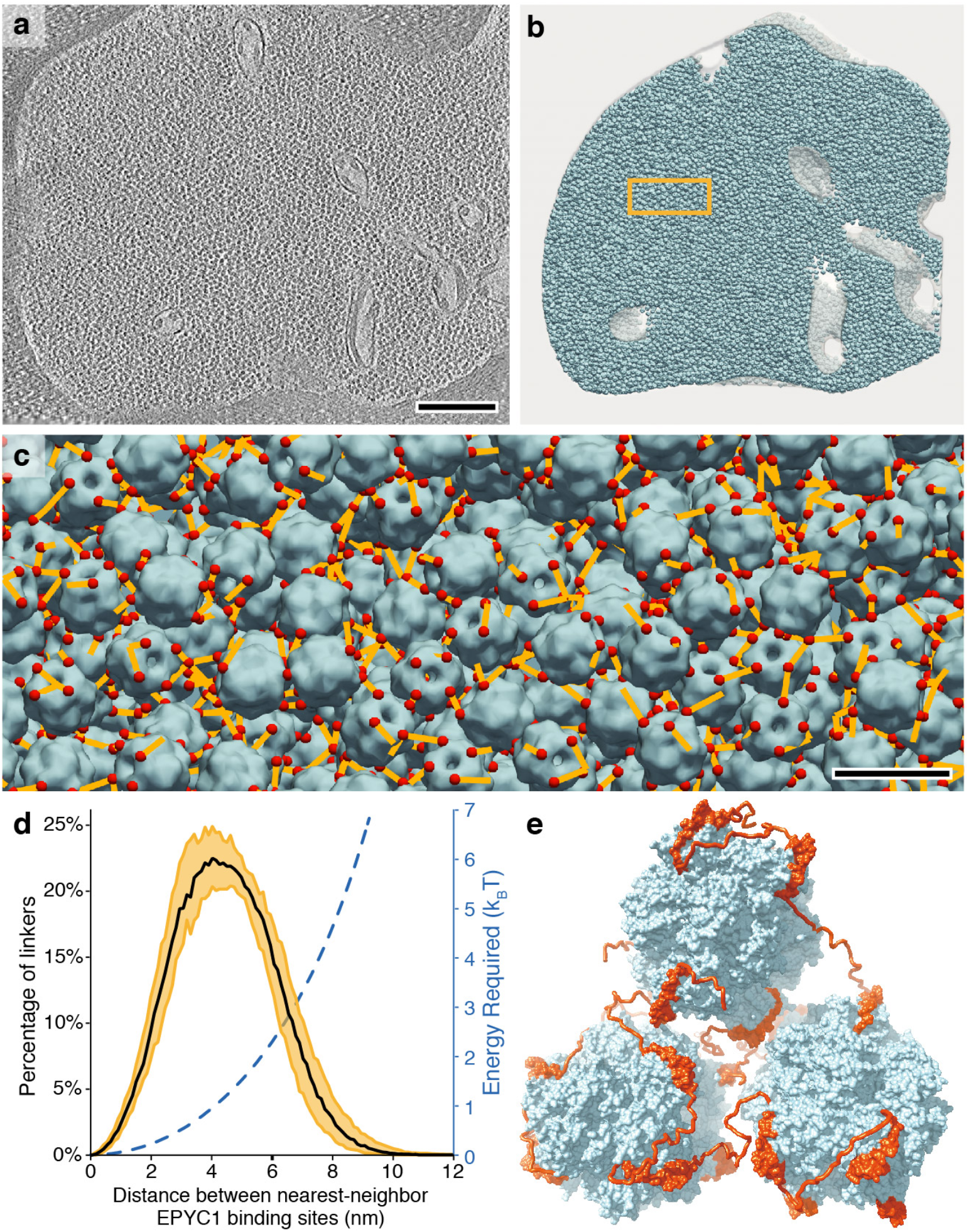
A model for matrix structure consistent with *in situ* Rubisco positions and orientations. **a**, The pyrenoid matrix was imaged by cryo-electron tomography^5^. An individual slice through the three-dimensional volume is shown. Scale bar = 200 nm. **b**, The positions and orientations of individual Rubisco holoenzymes (blue) were determined by subtomogram averaging and fit into the tomogram volume. **c**, The distances (yellow) between the nearest EPYC1-binding sites (red) on neighboring Rubisco holoenzymes (blue) were measured. The view is from inside the matrix; in some cases the nearest EPYC1 binding site is on a Rubisco that is out of the field of view, causing some yellow lines to appear unconnected in this image. Scale bar = 20 nm. **d**, Histogram showing the distances between the nearest EPYC1 binding sites on neighboring Rubisco holoenzymes. Shading indicates 95% confidence interval based on data from five independent tomograms. The estimated energy required for stretching a chain of 40 amino acids a given distance is shown in blue. **e**, A 3D model illustrates how EPYC1 (red) could crosslink multiple Rubisco holoenzymes (blue) to form the pyrenoid matrix. The conformations of the intrinsically disordered linkers between EPYC1 binding sites were modeled hypothetically.

A “linker” region of 40 amino acids is unlikely to be stretched to its maximum possible length of 14 nm *in vivo* due to the high entropic cost of this configuration. To determine whether a linker region can span the observed distances between nearest neighbor binding sites on adjacent Rubisco holoenzymes, we used a simple physics model to calculate the energy required to stretch a 40 amino acid chain to any given distance (Fig. 6d; see Methods). The model indicates that stretching the chain to ~7 nm requires an energy of 3 *k*_B_*T* (where *k*_B_, is the Boltzmann constant and *T* is the temperature), which could reasonably be borrowed from thermal fluctuations. Thus, our data suggest that the linker region between consecutive Rubisco-binding sites on the EPYC1 polypeptide can span the distance between adjacent Rubisco holoenzymes to hold the pyrenoid matrix together *in vivo*. It also appears likely that, in addition to bridging neighboring Rubisco holoenzymes, consecutive Rubisco-binding regions on a given EPYC1 may bind multiple sites on one Rubisco holoenzyme, as the distance between the nearest binding sites on the same holoenzyme is < 9 nm.

## Discussion

### Our data explain the structural basis of Rubisco condensation into a pyrenoid matrix

In this study, we have determined the structural basis for pyrenoid matrix formation for the first time in any species. We found that in the model alga Chlamydomonas, the intrinsically disordered protein EPYC1 has five regions of similar sequence that can bind to Rubisco as short peptides. These EPYC1 regions form an α-helix that binds to the Rubisco small subunit α-helices via salt bridges and hydrophobic interactions. EPYC1’s Rubisco-binding regions are spaced by linker sequences that are sufficiently long to span the distance between binding sites on adjacent Rubisco holoenzymes within the pyrenoid, allowing EPYC1 to serve as a molecular glue that clusters Rubisco together to form the pyrenoid matrix (Fig. 6e).

The multivalency of EPYC1 and the high K_D_ (~3 mM; Extended Data Fig. 1e) of individual Rubisco-binding regions are consistent with the emerging principle that cellular phase separation is mediated by weak multivalent interactions^20^. The high dissociation rate constant (>1/s; Extended Data Fig. 1d) of individual Rubisco-binding regions explains how the pyrenoid matrix can mix internally on the time scale of seconds^5^ despite the multivalency of EPYC1. The even spacing of the five Rubisco-binding regions across EPYC1 is noteworthy and may be an indication of selective pressure for an optimal distance between binding regions, and thus of an optimal spacing between Rubisco holoenzymes in the matrix.

Knowledge of the EPYC1-Rubisco binding mechanism now opens doors to the molecular characterization of the regulation of this interaction, which may govern the dissolution and condensation of the matrix during cell division^5^ and in response to environmental factors^21^. For example, phosphorylation of EPYC1^22^ may provide a mechanism to rapidly change the binding affinity of EPYC1 to Rubisco. Inactivation of one of EPYC1’s five Rubisco-binding regions would yield four binding regions, which would allow two EPYC1 molecules to form a mutually satisfied complex with each Rubisco, a configuration that is predicted to favor dissolution of the matrix^5^.

### The Rubisco-EPYC1 structure explains how other key pyrenoid proteins bind to Rubisco

In a parallel study (Meyer et al., please see unpublished manuscript provided as reference material), we recently discovered that a common sequence motif is present on many pyrenoid-localized proteins. The motif binds Rubisco, enabling recruitment of motif-containing proteins to the pyrenoid and mediating adhesion between the matrix, pyrenoid tubules, and starch sheath. This motif, [D/N]W[R/K]XX[L/I/V/A], is serendipitously present in EPYC1’s Rubisco-binding regions, and the motif residues mediate key binding interactions with Rubisco. In our structure, the R/K of the motif is represented by R64 of EPYC1, which forms a salt bridge with E24 of the Rubisco small subunit. The XX of the motif almost always includes a D or E; in our structure this feature is represented by E66 of EPYC1, which forms a salt bridge with R91 of the Rubisco small subunit. Finally, the W and [L/I/V/A] of the motif are represented by W63 and L67 of EPYC1, which contribute to the hydrophobic interactions with M87, L90 and V94 of the Rubisco small subunit. The key roles of the motif residues in the interface presented here strongly suggest that the structure we have obtained for one motif from EPYC1 also explains where and how all other variants of the motif, including those found on the key pyrenoid proteins SAGA1, SAGA2, RBMP1, RBMP2 and CSP41A, bind to Rubisco. Our observation that the Rubisco small subunit D23A/E24A and M87D/V94D mutants exhibit a more severe disruption of the pyrenoid than the *epyc1* mutant^10^ supports the idea that this region of Rubisco interacts not only with EPYC1, but also with other proteins required for pyrenoid biogenesis, making this binding interaction a central hub of pyrenoid biogenesis.

### There are structural similarities and differences between the pyrenoid matrix and bacterial carboxysomes

Although α- and β-carboxysomes are morphologically, functionally and evolutionarily distinct from the pyrenoid, their Rubisco is also thought to be clustered by linker proteins. Like EPYC1, the α-carboxysome linker protein CsoS2^23^ is intrinsically disordered and is proposed to bind Rubisco as an unfolded peptide^24^. In contrast, the β-carboxysome linker protein CcmM has been proposed to bind Rubisco using folded globular domains^25,26^. The use of an unfolded peptide as in the case of EPYC1 and CsoS2 may provide the benefit of requiring fewer amino acids for achieving the desired binding function. A notable difference is the location of the binding site on

Rubisco: whereas both carboxysomal linker proteins bind to the interface between two Rubisco large subunits^24,26^, EPYC1 binds to the Rubisco small subunit. It remains to be seen whether this difference in binding site has functional consequences, such as impacts on the three-dimensional packing of Rubisco.

### Our findings advance the ability to engineer a pyrenoid into crops

There is currently significant interest in engineering Rubisco condensates into monocotyledonous crops such as wheat and rice to enhance yields^27–30^. Binding of EPYC1 to the Rubisco small subunit presents a promising route for engineering a Rubisco condensate, as the Rubisco small subunit is encoded in the nuclear genome, making it more easily amenable to genetic modification in those crops than the chloroplast-encoded Rubisco large subunit^31^. Knowledge of the binding mechanism now allows engineering of minimal sequence changes into native crop Rubiscos to enable binding to EPYC1 and to other key proteins required to reconstitute a functional pyrenoid.

### Our work provides insights into pyrenoid matrix formation in other species

Pyrenoids appear to have evolved independently in different lineages through convergent evolution^7,32^. EPYC1, its Rubisco-binding sequences, and the amino acid residues that form the EPYC1 binding site on the surface of Rubisco are conserved across the order Volvocales, as evidenced from the genome sequences of *Tetrabaena socialis, Gonium pectorale* and *Volvox carteri* (Extended Data Table 2). While the molecular mechanisms of matrix formation in other lineages remain to be uncovered, candidate linker proteins have been identified based on similarity of sequence properties to EPYC1^10^. We hypothesize that the matrix in other lineages is formed based on similar principles to those we observed in Chlamydomonas. Our experimental approach for characterizing the binding interaction provides a roadmap for future structural studies of pyrenoids across the tree of life.

### This study provides a high-resolution structural view of a phase separated organelle

The pyrenoid matrix presents an unusual opportunity to study a two-component molecular condensate where one of the components, Rubisco, is large and rigid, and the other component, EPYC1, is a simple intrinsically disordered protein consisting of nearly identical tandem repeats. The rigidity and size of Rubisco holoenzymes previously enabled the determination of their positions and orientations within the pyrenoid matrix of native cells by cryo-electron tomography^5^. The identification of EPYC1 binding sites on Rubisco in the present work and the modeling of linker regions between EPYC1’s Rubisco binding regions now make the Chlamydomonas pyrenoid matrix one of the most structurally well-defined phase separated organelles. Thus, beyond advancing our structural understanding of pyrenoids, organelles that play a central role in the global carbon cycle, we hope that the findings presented here will also more broadly enable advances in the biophysical understanding of phase separated organelles.

## Methods

### Strains and culture conditions

Chlamydomonas wild-type (WT) strain cMJ030 was maintained in the dark or low light (~10 μmol photons m^−2^ s^−1^) on 1.5% agar plates containing Tris-Acetate-Phosphate medium with revised trace elements^33^. For Rubisco extraction, 500 mL Tris-Acetate-Phosphate medium in a 1 L flask was inoculated with a loopful of cells and the culture was grown to 4 x 10^6^ cells/mL at 22°C, shaking at 200 rpm under ~100 μmol photons m^−2^ s^−1^ white light in 3% CO_2_. Chlamydomonas mutant T60-3^34^ (Δ*rbcs*; containing a deletion of both *RBCS* genes) was used for generating Rubisco small subunit point mutants and a wild-type control in the same background. This strain was maintained on agar in the dark or low light (~10 μmol photons m^−2^s^−1^).

### Protein extraction

Rubisco was purified from Chlamydomonas strain cMJ030^35^. Cells were disrupted by ultrasonication in lysis buffer (10 mM MgCl_2_, 50 mM Bicine, 10 mM NaHCO_3_, 1 mM dithiothreitol, pH 8.0) supplemented with Halt Protease Inhibitor Cocktail, EDTA-Free (Thermo Fisher Scientific). The soluble lysate was fractionated by ultracentrifugation on a 10-30% sucrose gradient in a SW 41 Ti rotor at a speed of 35,000 rpm for 20 hours at 4°C. Rubisco-containing fractions were applied to an anion exchange column (MONO Q 5/50 GL, GE Healthcare) and eluted with a linear salt gradient from 30 to 500 mM NaCl in lysis buffer.

### Peptide arrays

Peptide arrays were purchased from the MIT Biopolymers Laboratory (Cambridge, MA). The tiling array was composed of 18-amino-acid peptides that tiled across the full-length EPYC1 sequence with a step size of one amino acid. Each peptide was represented by at least two spots on the array, and these replicates were averaged during data analysis. The locations of peptides on the array were randomized. In the substitution arrays, peptides were designed to represent every possible one-amino-acid mutation of the indicated region on EPYC1 by substitution with one of the other 19 amino acids. The arrays were activated by methanol, then washed 3×10 min in binding buffer (50 mM HEPES, 50 mM KOAc, 2 mM Mg(OAc)_2_.4H_2_O, 1 mM CaCl_2_ and 200 mM sorbitol, pH 6.8). The arrays were then incubated at 4°C with 1 mg purified Rubisco overnight. The arrays were washed in binding buffer to remove any unbound Rubisco. Using a semi-dry transfer apparatus (BIO-RAD), bound Rubisco was transferred onto an Immobilon-P PVDF membrane (Millipore Sigma). Rubisco was immuno-detected with a polyclonal primary antibody raised against Rubisco^15^ (1:10,000) followed by a HRP conjugated goat anti-rabbit (1:20,000; Invitrogen). Arrays were stripped with Restore™ Western Blot Stripping Buffer before re-use (Thermo Fisher Scientific).

### Surface plasmon resonance (SPR) experiments

All the surface preparation experiments were performed at 25°C using a Biacore 3000 instrument (GE Healthcare). Purified Rubisco was immobilized on CM5 sensor chips using a Biacore Amine Coupling Kit according to the manufacturer’s instructions. Briefly, the chip surface was activated by an injection of 1:1 N-hydroxysuccinimide (NHS)/1-ethyl-3-(3-dimethylaminopropyl)carbodiimide hydrochloride (EDC). Rubisco was diluted to ~100 μg/mL in 10 mM acetate (pH 4.5; this pH had been previously optimized using the immobilization pH scouting wizard) and was injected over the chip surface. Excess free amine groups were then capped with an injection of 1 M ethanolamine. Typical immobilization levels were 8,000 to 10,000 resonance units (RU), as recommended for binding experiments of small molecules. For kinetic experiments (for determining the binding affinities), the typical immobilization levels were ~5,000 RU. The control surfaces were prepared in exactly the same manner as the experimental surfaces except that no Rubisco was injected. For immobilizations, the running buffer was the Biacore HBS-EP Buffer (0.01 M HEPES pH 7.4, 0.15 M NaCl, 3 mM EDTA, 0.005% v/v Surfactant P20).

All the binding assays were performed using the Biacore PBS-P+ Buffer (20 mM phosphate buffer, 2.7 mM KCl, 137 mM NaCl and 0.05% Surfactant P20, pH 6.8) as a running buffer, as recommended for small molecule analysis in Biacore systems. The analytes, consisting of EPYC1 peptides synthesized by Genscript (Piscataway, New Jersey), were dissolved in the same running buffer and diluted to 1 mM. The analytes were injected over the control surface and experimental surfaces at a flow rate of 26 μL/min for 2.5 minutes, followed by 2.5 minutes of the running buffer alone to allow for dissociation. The surfaces were then regenerated using running buffer at a flow rate of 30 μL/min for 10 minutes. In all cases, binding to the control surface was negligible.

For determining the K_D_ of EPYC1 peptide, the kinetic assays were performed with a running buffer consisting of 200 mM sorbitol, 50 mM HEPES, 50 mM KOAc, 2 mM Mg(OAc)_2_.4H_2_O and 1 mM CaCl_2_ at pH 6.8 (the same buffer as the peptide array assay). The EPYC1 peptide was dissolved in the same running buffer as the assay and the serial dilutions were also made in the same buffer. The analytes were injected over the control surface and experimental surfaces at a flow rate of 15 μL/min for 2 minutes, followed by 10 minutes with the running buffer alone to allow for dissociation. The surfaces were then regenerated by the running buffer at a flow rate of 30 μL/min for 10 minutes. In all cases, binding to the blank chip was negligible. The fitting and modeling were performed with the BIAevaluation software.

### Single-particle cryo-electron microscopy data collection and image processing

Rubisco and peptide with the final concentration of 1.69mg/ml (=3.02 μM) and 7.5 mM were incubated on ice for 20 minutes in buffer consisting of 200 mM sorbitol, 50 mM HEPES, 50 mM KOAc, 2 mM Mg(OAc)_2_.4H_2_O and 1 mM CaCl_2_ at pH 6.8 (the same buffer as the peptide array assay and the SPR binding assay). For both apo Rubisco and Rubisco incubated with peptide, similar cryo grid-making procedures were used. 400-mesh Quantifoil 1.2/1.3 Cu grids (Quantifoil, Großlöbichau, Germany) were made hydrophilic by glow discharging for 60 seconds with a current of 15 mA in a Pelico EasiGlow system. Samples on cryo grids were plunge-frozen using an FEI Mark IV Vitrobot (FEI company, part of Thermo Fisher Scientific, Hillsboro, OR). The chamber of the Vitrobot was kept at 4°C and 100% relative humidity. 3 μl of sample was applied to the glow-discharged grid, blotted with filter paper for 3 seconds with the equipment-specific blotting force set at 3. After blotting, the grid was rapidly plunge-frozen into a liquid ethane bath.

Cryo grids were loaded into a 300 kV FEI Titan Krios cryo electron microscope (FEI Company) at HHMI Janelia Research Campus, Janelia Krios2, equipped with a Gatan K2 Summit camera. After initial screening and evaluation, fully automated data collection was carried out using SerialEM. The final exposure from each collection target was collected as a movie utilizing dose fractionation on the K2 Summit camera operated in super-resolution mode. The movie was collected at a calibrated magnification of 38,168x, corresponding to 1.31 Å per physical pixel in the image (0.655 Å per super-resolution pixel). The dose rate on the specimen was set to be 5.82 electrons per Å^2^ per second and total exposure time was 10 s, resulting in a total dose of 58.2 electrons per Å^2^. With dose fractionation set at 0.2 s per frame, each movie series contained 50 frames and each frame received a dose of 1.16 electrons per Å^2^. The spherical aberration constant of the objective lens is 2.7 mm and an objective aperture of 100 μm was used. The nominal defocus range for the automated data collection was set to be between −1.5 μm and −3.0 μm.

The movies were 2x binned and motion corrected using MotionCor2^36^ and CTF was estimated using CTFFIND^37^ in Relion 3.0^38^. The particles were selected using cisTEM^39^ and 1,809,869 peptide bound Rubisco particles and 677,071 particles in the apo state were extracted with a box size of 192×192pixels. 2D classification was performed using cisTEM 2D. The classes presenting detailed features in class averages were chosen for 3D classification on cryoSPARC^40,41^ for peptide-bound Rubisco and on Relion for the apo state. The 3D class showing clear secondary structures was chosen for 3D auto-refine first without symmetry and then with D4 symmetry imposed. After CTF refinement and Bayesian polishing in Relion, the reconstructed map resolution is 2.68 Å for the apo state and 2.62 Å for the peptide bound state. Details for single-particle cryo-EM data collection and image processing are included in the Extended Data Table 1.

### Single-particle cryo-electron microscopy model building, fitting, and refinement

A full model for Rubisco from Chlamydomonas was produced from an X-ray structure^13^ (PDB entry 1GK8) and used for rigid body fitting into a local resolution filtered cryo-EM map with an average resolution of 2.62 Å using UCSF Chimera^42^. After rigid body fitting of the full complex, initial flexible fitting was performed in COOT^43^ by manually going through the entire peptide chain of a single large and small Rubisco subunit before applying the changes to the other seven large and small subunits. The sequence of the peptide was used to predict secondary structure elements using JPred4^44^ which resulted in the prediction that the C-terminal region (NWRQELES) is α-helical. Guided by this prediction, the peptide was built manually into the density using COOT. Additional real space refinement of the entire complex was performed using Phenix^45^. Models were subjected to an all-atom structure validation using MolProbity^46^. Figures were produced using UCSF Chimera.

### Liquid–liquid phase separation assay

Proteins used in the liquid–liquid phase separation assay were obtained and stored essentially as described previously^11^. Briefly, Rubisco was purified from *C. reinhardtii* cells (CC-2677 cw15 nit1-305 mt-5D, Chlamydomonas Resource Center) grown in Sueoka’s high-salt medium^47^, using a combination of anion exchange chromatography and gel filtration.

The EPYC1 full-length gene (encoding amino acids 1-317) and corresponding R/K mutant (EPYC1^R64A/K127A/K187A/K248A/R314A^) were synthesized by GenScript and cloned between the SacII and HindIII site of the pHue vector^48^. Proteins were produced in the *E. coli* strain BL21 (DE3) harbouring pBADESL^49^ for co-expression of the *E. coli* chaperonin GroEL/S. The purification was conducted with minor changes (dialysis for removal of high immidazol concentrations was skipped by running the gel-filtration column before the second IMAC). After the first IMAC step and cleavage^50^ of the N-terminal His_6_–ubiquitin tag, proteins were separated by gel filtration. Finally, the peak fraction was passed a second time through an IMAC column, collecting EPYC1 from the flow through.

EPYC1-Rubisco condensates were reconstituted *in vitro* in a buffer containing 20 mM Tris-HCl (pH 8.0) and NaCl concentrations as indicated. 5 μl reactions were incubated for 3 min at room temperature before monitoring the droplet formation by differential interference contrast (DIC) microscopy. DIC images were acquired with a Nikon Eclipse Ti Inverted Microscope using a 60 × oil-immersion objective after allowing the droplets to settle on the coverslip (Superior Marienfeld, Germany) surface for about 3 min. For droplet sedimentation assays 10 μl reactions were incubated for 3 min at 20°C before separating the droplets form the bulk phase by spinning for 3 min at 21,000xg and 4°C. Pelleted droplets and supernatant fractions were analyzed using Coomassie-stained SDS-PAGE.

### Yeast two-hybrid assay

Yeast two-hybrid to detect interactions between EPYC1 and RbcS1 was carried out as described previously^12^. EPYC1 was cloned into the two-hybrid vector pGBKT7 to create a fusion with the GAL4 DNA binding domain. Point mutations were introduced by PCR into RbcS1, which was then cloned in the pGADT7 to create a fusion with the GAL4 activation domain. Yeast cells were then co-transformed with binding and activation domain vectors. Successful transformants were cultured, diluted to an optical density at 600 nm (OD600) of 0.5 or 0.1, and plated onto SD-L-W and SD-L-W-H containing increasing concentrations of the HIS3 inhibitor triaminotriazole (3-AT). Plates were imaged after 3 days. Spots shown in Fig. 5a were grown at 5 mM 3-AT from a starting OD600 of 0.5; they are a subset of the full dataset shown in Extended Data Fig. 6.

### Cloning of Rubisco small subunit point mutants

The plasmid pSS1-ITP^51^ which contains Chlamydomonas *RBCS1* including UTRs and introns 1 and 2 was used as a starting point for generating plasmids pSH001 and pSH002, which encode RBCS1^D23A/E24A^, and RBCS1^M87D/V94D^, respectively. The point mutations were generated by Gibson assembly^52^ of gBlocks (synthesized by Integrated DNA Technologies) containing the desired mutations into pSS-ITP that had been enzyme digested by restriction endonucleases (XcmI and BbvCI for the D23A/E24A mutations and BbvCI and BlpI for the M87D/V94D mutations). All constructs were verified by Sanger sequencing.

The fragment for making pSH001 (containing the D23A/E24A Rubisco small subunit mutant) had the following sequence:

GCAGGGCTGCCCCGGCTCAGGCCAACCAGATGATGGTCTGGACCCCGGTCAACAACAAGATGTTCGAGACCTTCTCCTACCTGCCTCCTCTGACCGCCGCGCAGATCGCCGCCCAGGTCGACTACATCGTCGCCAACGGCTGGATCCCCTGCCTGGAGTTCGCTGAGGCCGACAAGGCCTACGTGTCCAAC

The fragment for making pSH002 (containing the M87D/V94D Rubisco small subunit mutant) had the following sequence:

CTGCCTGGAGTTCGCTGAGGCCGACAAGGCCTACGTGTCCAACGAGTCGGCCATCCGCTTCGGCAGCGTGTCTTGCCTGTACTACGACAACCGCTACTGGACCATGTGGAAGCTGCCCATGTTCGGCTGCCGCGACCCCGACCAGGTGCTGCGCGAGATCGACGCCTGCACCAAGGCCTTCCCCGATGCCTACGTGCGCCTGGTGGCCTTCGACAACCAGAAGCAGGTGCAGATCATGGGCTTCCTGGTCCAGCGCCCCAAGACTGCCCGCGACTTCCAGCCCGCCAACAAGCGCTCCGTGTAAATGGAGGCGCTCGTCGATCTGAGCCGTGTGTGATGTTTGTTGGTGTTTGAGCGAGTGCAATGAGAGTGTGTGTGTGTGTGTTGTTGGTGTGTGGCTAAGCCAAGCGTGATCGC

### Transformation of Chlamydomonas to make the Rubisco small subunit point mutants

Chlamydomonas strains *Δrbcs;RBCS^WT^, Δrbcs;RBCS^D23A/E24A^*, and *Δrbcs;RBCS^M87D/V94D^* were generated by transforming pSS1-ITP, pSH001, and pSH002 (encoding Rubisco small subunit constructs) into the Rubisco small subunit deletion mutant T60 (*Δrbcs*) by electroporation as described previously^53^. For each transformation, 29 ng kbp^−1^ of KpnI linearized plasmid was mixed with 250 μL of 2 x 10^8^ cells mL^−1^ at 16°C and electroporated immediately. Transformant colonies were selected on Tris-Phosphate plates without antibiotics at 3% v/v CO_2_ under ~50 μmol photons m^−2^ s^−1^ light. The sequence of RbcS in the transformants was verified by PCR amplification and Sanger sequencing.

### Spot tests

*Δrbcs;RBCS^WT^, Δrbcs;RBCS^D23A/E24A^*, and *Δrbcs;RBCS^M87D/V94D^* were grown in Tris-Phosphate medium at 3% CO_2_ until ~2×10^6^ cells mL^−1^. Cells were diluted in Tris-Phosphate medium to a concentration of 8.7 x10^7^ cells mL^−1^, then serially diluted 1:10 three times. 7.5 μL of each dilution was spotted onto four TP plates and incubated in air or 3% CO_2_ under 20 or 100 μmol photons m^−2^ s^−1^ white light for 9 days before imaging.

### Transmission electron microscopy

Samples for electron microscopy were fixed for 1 hour at room temperature in 2.5% glutaraldehyde in Tris-Phosphate medium (pH 7.4), followed by 1 hour at room temperature in 1% OsO_4_, 1.5% K_3_Fe(CN)_3_, and 2 mM CaCl_2_. Fixed cells were then bulk stained for 1 hour in 2% uranyl acetate, 0.05 M maleate buffer at pH 5.5. After serial dehydration (50%, 75%, 95%, and 100% ethanol, followed by 100% acetonitrile), samples were embedded in epoxy resin containing 34% Quetol 651, 44% nonenyl succinic anhydride, 20% methyl-5-norbornene-2,3-dicarboxylic anhydride, and 2% catalyst dimethylbenzylamine. Ultramicrotomy was done by the Core Imaging Lab, Medical School, Rutgers University. Imaging was performed at the Imaging and Analysis Center, Princeton University, on a CM100 transmission electron microscope (Philips, Netherlands) at 80 kV.

### Measurement of nearest-neighbor distances between EPYC1-binding sites on Rubisco holoenzymes within pyrenoids

For detailed descriptions of the Chlamydomonas cell culture, vitrification of cells onto EM grids, thinning of cells by cryo-focused ion beam milling, 3D imaging of native pyrenoids by cryo-electron tomography, tomographic reconstruction, and subtomogram averaging, see our previous study^5^. In that study, we measured the distances between the center positions of Rubisco complexes within tomograms of five pyrenoids. The spatial parameters determined in that study were combined with the EPYC1-binding sites resolved here by cryo-EM single-particle analysis to measure the nearest-neighbor distances between EPCY1-binding sites on adjacent Rubisco complexes within the native pyrenoid matrix.

The *in situ* subtomogram average EMD-3694^5^ was used as the reference for the Rubisco model. We extracted the isosurface from this density using the 0.5 contour level recommended in the Electron Microscopy Data Bank entry. We then fit the atomic model of EPYC1-bound Rubisco (Fig. 2) within the EMD-3694 density, and for each EPYC1-binding site, we marked the closest point on the isosurface to define the EPYC1 binding sites on this model. The positions and orientations previously determined by subtomogram averaging were used to place each Rubisco model and its corresponding binding sites into the pyrenoid tomograms using the PySeg program^54^.

To compute the nearest-neighbor distances between EPYC1-binding sites on two different Rubisco complexes, first, linkers were drawn between each EPYC1 binding site and all other binding sites within 25 nm. Binding sites on the same Rubisco complex were ignored. Next, the linkers were filtered by length (defined as the Euclidean distance between the two binding sites), and only the shortest linker was retained for each binding site. To prevent edge effects, linkers were discarded if they had a binding site <12 nm from the masked excluded volume (grey in Fig. 6b), which marks the border of the analyzed pyrenoid matrix. Finally, linker distances were plotted in a histogram to show the distribution of lengths (normalized to 100%).

### Modeling of the energy required to stretch EPYC1-linker regions

The energy required to stretch the linker regions between EPYC1’s Rubisco-binding regions was determined as follows. The force F required to stretch a 40 amino acid linker region to any given length z was approximated using a wormlike chain model^55^:

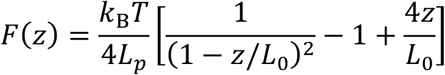

In the above equation, *k*_B_ is the Boltzmann constant, *T* is the temperature, L_p_ is the persistence length (assumed to be 1 nm, a representative value for disordered proteins), and L_0_ is the contour length (estimated as 40 amino acids * 0.36 nm/amino acid). The energy required to stretch the linker to a length x is given by:

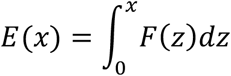

This energy was calculated and plotted in Fig. 6d.

## Acknowledgements

We thank Jianping Wu, Nieng Yan, Luke Mackinder, Cliff Brangwynne and members of the Jonikas laboratory for helpful discussions; Ned Wingreen, Silvia Ramundo, Jessi Hennacy, and Eric Franklin for constructive feedback on the manuscript; Wolfgang Baumeister and Jürgen Plitzko for providing support and cryo-ET instrumentation; and Miroslava Schaffer for help with acquiring the cryo-ET data, previously published in Freeman Rosenzweig et al., 2017. This project was funded by National Science Foundation (IOS-1359682 and MCB-1935444), National Institutes of Health (DP2-GM-119137), and Simons Foundation and Howard Hughes Medical Institute (55108535) grants to M.C.J., Deutsche Forschungsgemeinschaft grant (EN 1194/1-1 as part of FOR2092) to B.D.E., Ministry of Education (MOE Singapore) Tier 2 grant (MOE2018-T2-2-059) to O.M.-C., UK Biotechnology and Biological Sciences Research Council (BB/S015531/1) and Leverhulme Trust (RPG-2017-402) grants to A.J.M and N.A., NIH grant R01GM071574 to F.M.H., Deutsche Forschungsgemeinschaft fellowship (PO2195/1-1) to S.A.P., and National Institute of General Medical Sciences of the National Institutes of Health (T32GM007276) training grant to V.K.C.. The content is solely the responsibility of the authors and does not necessarily represent the official view of the National Institutes of Health.

## Author contributions

S.H., P.D.J., V.C., F.M.H., T.W., O.M.-C., B.D.E., and M.C.J. designed experiments. S.H. identified EPYC1’s Rubisco-binding regions on EPYC1 by peptide tiling array and SPR. S.H. and S.A.P. prepared the Rubisco and EPYC1 peptide sample for single-particle cryo-EM; S.H., S.A.P. and G.H. prepared the Rubisco samples for peptide tiling array and surface plasmon resonance. H.-T.C., D.M. and Z.Y. performed Cryo-EM grid preparation, sample screening, data acquisition, image processing, reconstruction and map generation. D.M. and P.D.J. carried out single-particle model building and fitting and refinement. S.H., H.-T.C., D.M., P.D.J., F.M.H. and M.C.J. analyzed the structures. S.H. and W.P. analyzed EPYC1 binding to Rubisco by peptide substitution array and SPR. T.W. performed in vitro reconstitution phase separation experiments. N.A. and A.J.M. performed yeast two-hybrid experiments. S.H. and M.T.M. made Rubisco small subunit point mutants. S.H. performed spot test experiments. M.T.M. performed TEM. A.M.-S. performed the cryo-ET data analysis and modeling. S.H. and M.C.J. wrote the manuscript. All authors read and commented on the manuscript.

---

**Extended Data Fig. 1.**
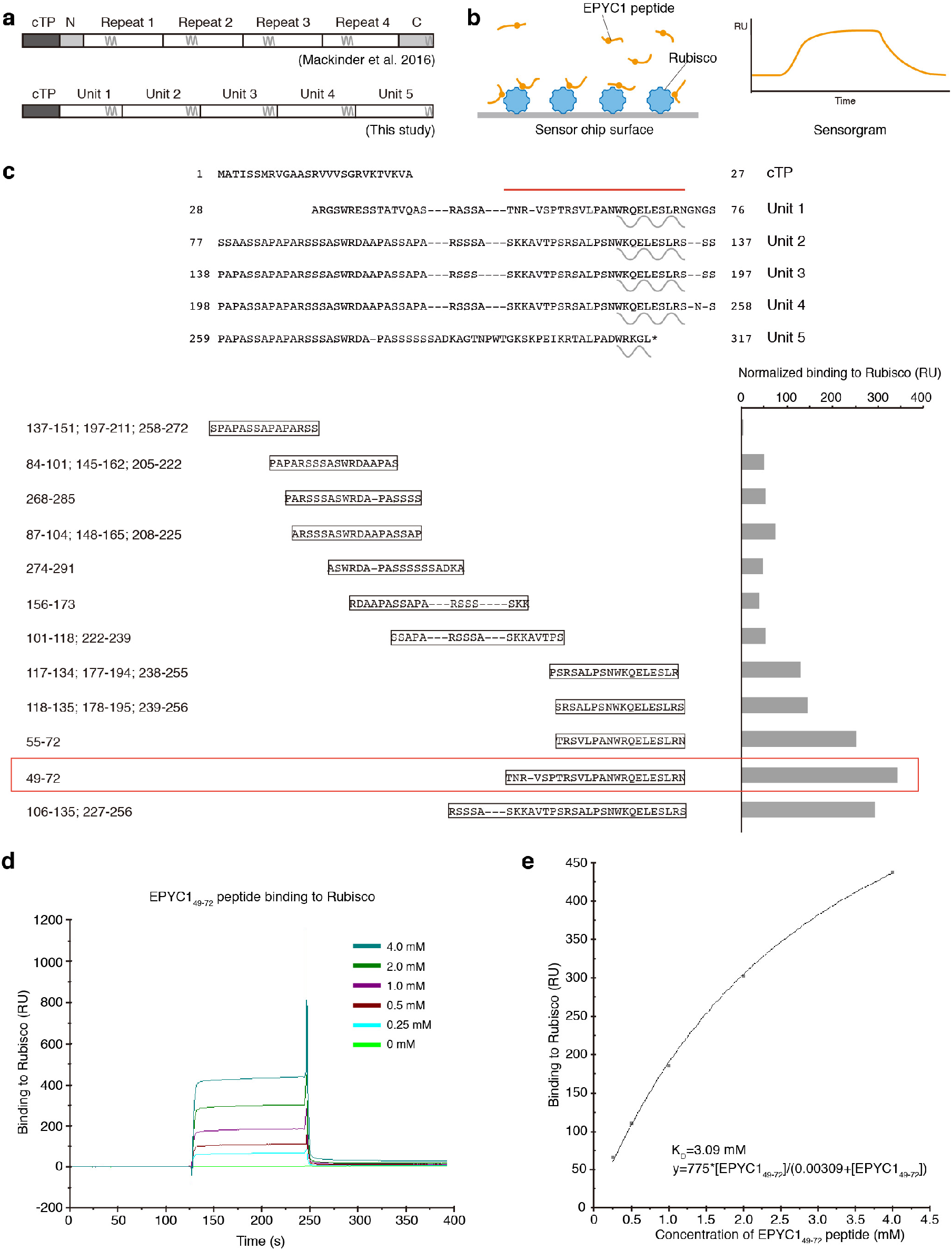
The EPYC1 peptide with the highest binding affinity to Rubisco was chosen for structural studies. **a**, Diagram indicating the differences between the previously defined sequence repeats^10^ and the newly defined sequence repeats on full-length EPYC1. **b**, To verify the Rubisco-binding regions on EPYC1, surface plasmon resonance (SPR) was used to measure the binding of EPYC1 peptides to Rubisco. Purified Rubisco was immobilized on a sensor surface, and the EPYC1 peptides in solution were injected over the surface. The binding activity was recorded in real time in a sensorgram. **c**, The peptides used in SPR experiments are shown aligned to the sequence as shown in Fig. 1. The Rubisco-binding signal from the SPR experiment of each peptide is shown after normalization to the peptide’s molecular weight. EPYC149-72 was chosen for structural studies based on its reproducible high Rubisco binding signal. **d**, The Rubisco-binding response of the EPYC149-72 peptide at different concentrations was measured by SPR. **e**, The binding responses shown in (d) were fitted to estimate the K_D_ of EPYC149-72 peptide binding to Rubisco.

**Extended Data Fig. 2.**
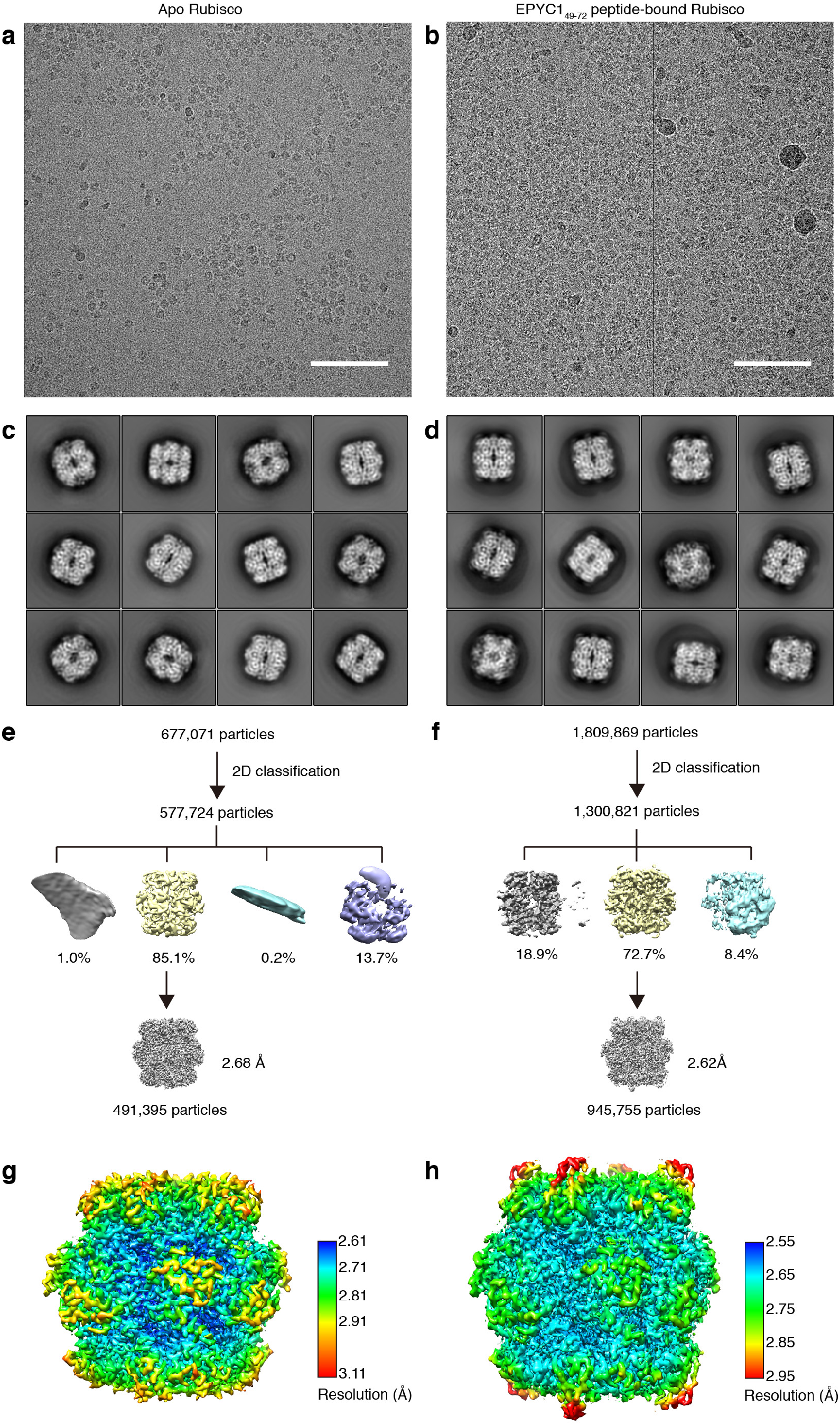
Cryo-EM data collection and image processing procedure. a, Representative micrograph of the apo Rubisco sample. Scale bar = 100 nm. b, Representative micrograph of Rubisco-EPYC149-72 complexes. Scale bar = 100 nm. c, Representative 2D class averages of the apo Rubisco sample. d, Representative 2D class averages of Rubisco-EPYC149-72 complexes. e, Overview of the workflow for single particle data processing for the apo Rubisco sample. f, Overview of the workflow for single particle data processing for the Rubisco-EPYC1_49-72_ sample. g, Local resolution estimation diagram of the final refined apo Rubisco map. **h**, Local resolution estimation diagram of the final refined Rubisco-EPYC1_49-72_ complexes map.

**Extended Data Fig. 3.**
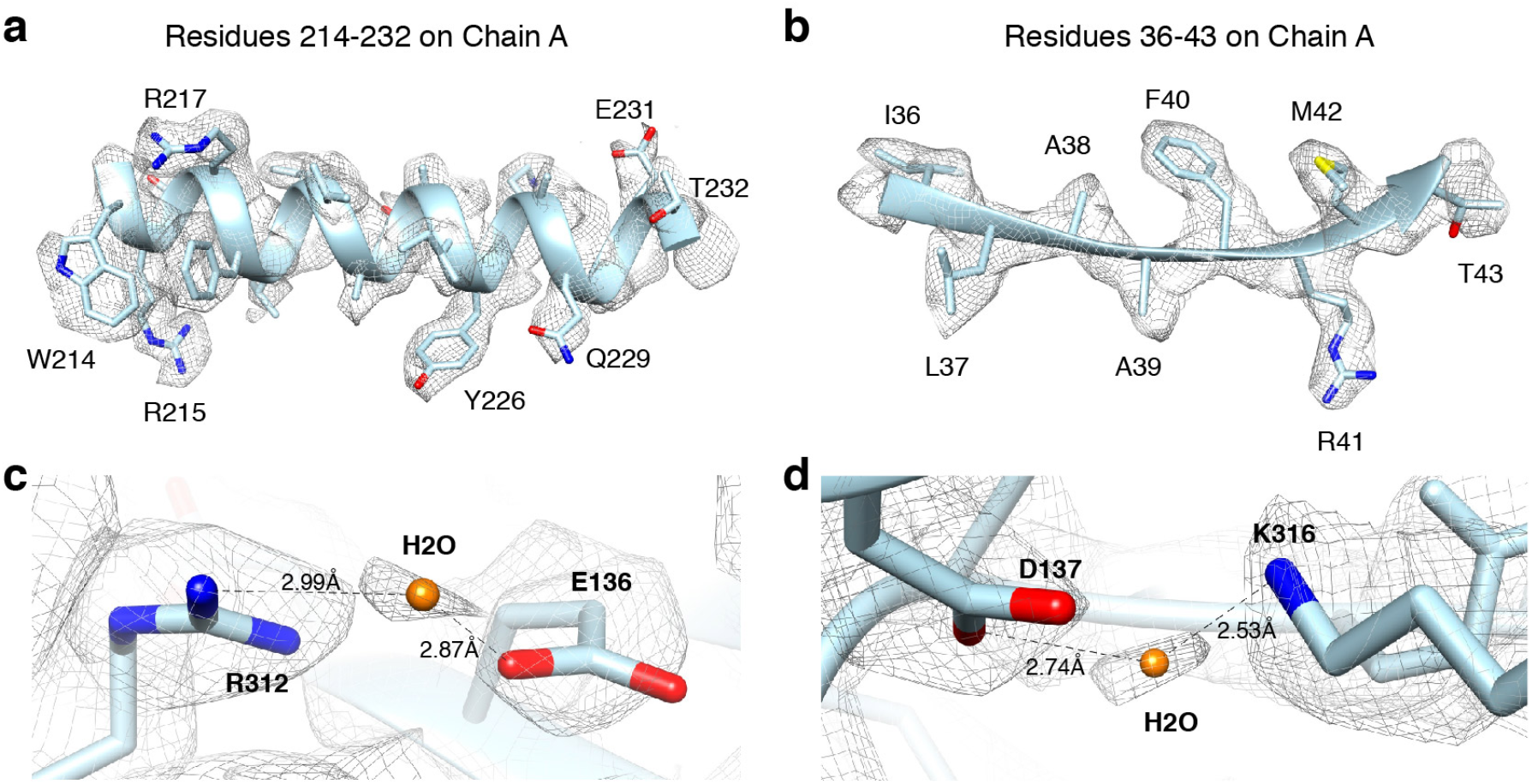
Cryo-EM analysis and resolution of Rubisco-EPYC1 peptide complexes in this study. **a-b,** Representative cryo-EM density quality showing an α-helix of residues 214-232 in chain A (a) (one of the Rubisco large subunits) and a ß-sheet of residues 36-43 in chain A (b). The densities are shown as meshwork in gray. The backbones of the structural model are in ribbon representation, and side chains are shown in stick representation. **c-d,** Representative cryo-EM density quality showing water molecules as orange spheres. One water molecule between R312 and E136 on chain A is shown in panel c, and another water molecule between D137 and K316 on chain A is shown in panel d.

**Extended Data Fig. 4.**
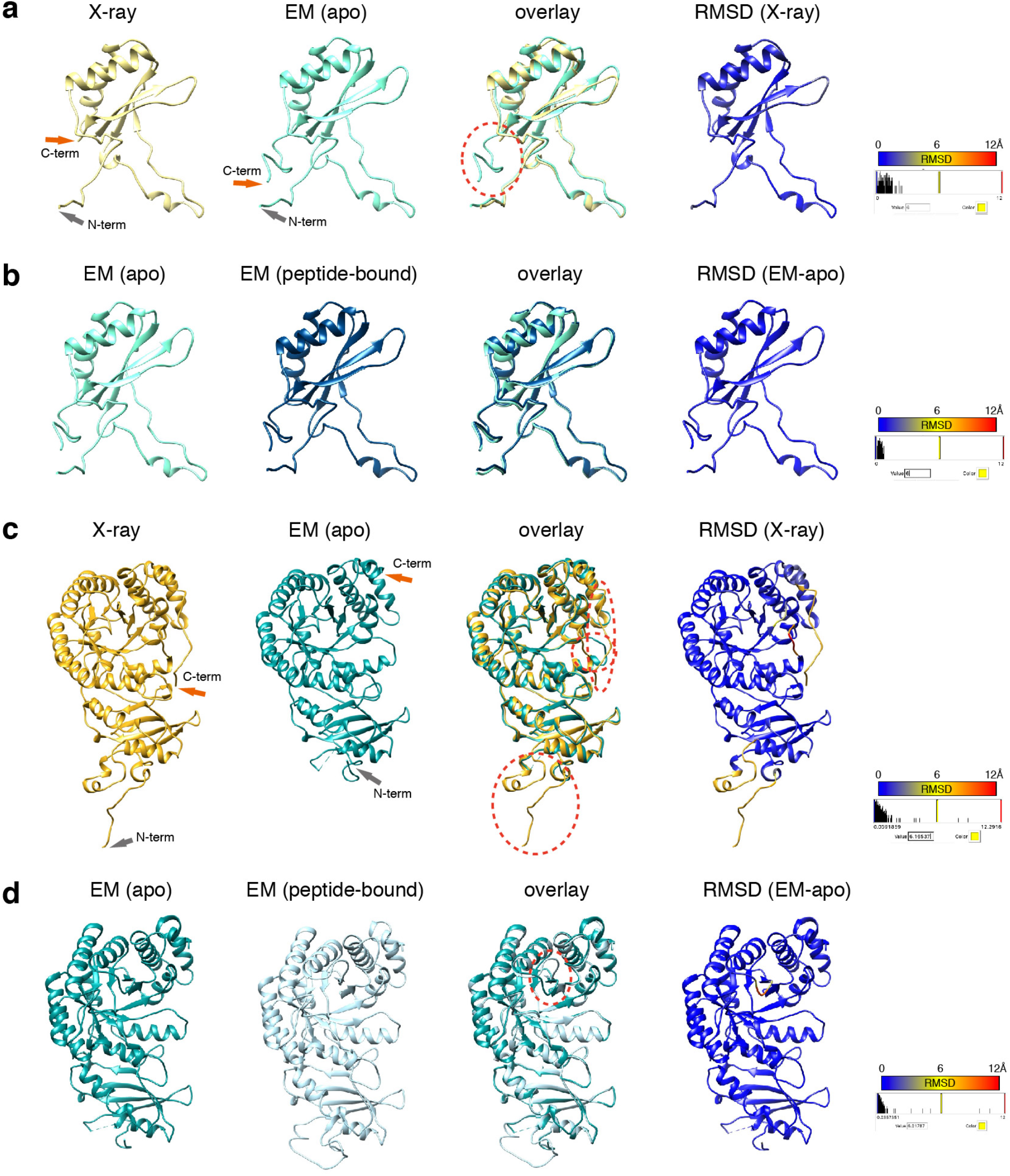
Comparison of our EM structure and the published X-ray crystallography structure (1gk8) of Rubisco purified from *Chlamydomonas reinhardtii^13^*, and comparison of our EM structure of native Rubisco and Rubisco bound with EPYC149-72 peptide. a, Comparison of the structure of the small subunit of apo Rubisco obtained here by EM with 1gk8. The EM structure has additional C-terminus density past residue 126, circled by a red dashed line. b, Comparison of our two EM structures of the small subunit: from apo Rubisco and from EPYC1 peptide-bound Rubisco. c, Comparison of the structure of the large subunit of apo Rubisco obtained here by EM with 1gk8. The three major differences found between the X-ray structure and the EM structure of the large subunit are circled with red dashed lines. d, Comparison of our two EM structures of the large subunit: from apo Rubisco and from EPYC1 peptide-bound Rubisco. The major difference found between the peptide-bound structure and the apo EM structure was the loop between K175 and L180 of the large subunit, which is shown circled by a red dashed line.

**Extended Data Fig. 5.**
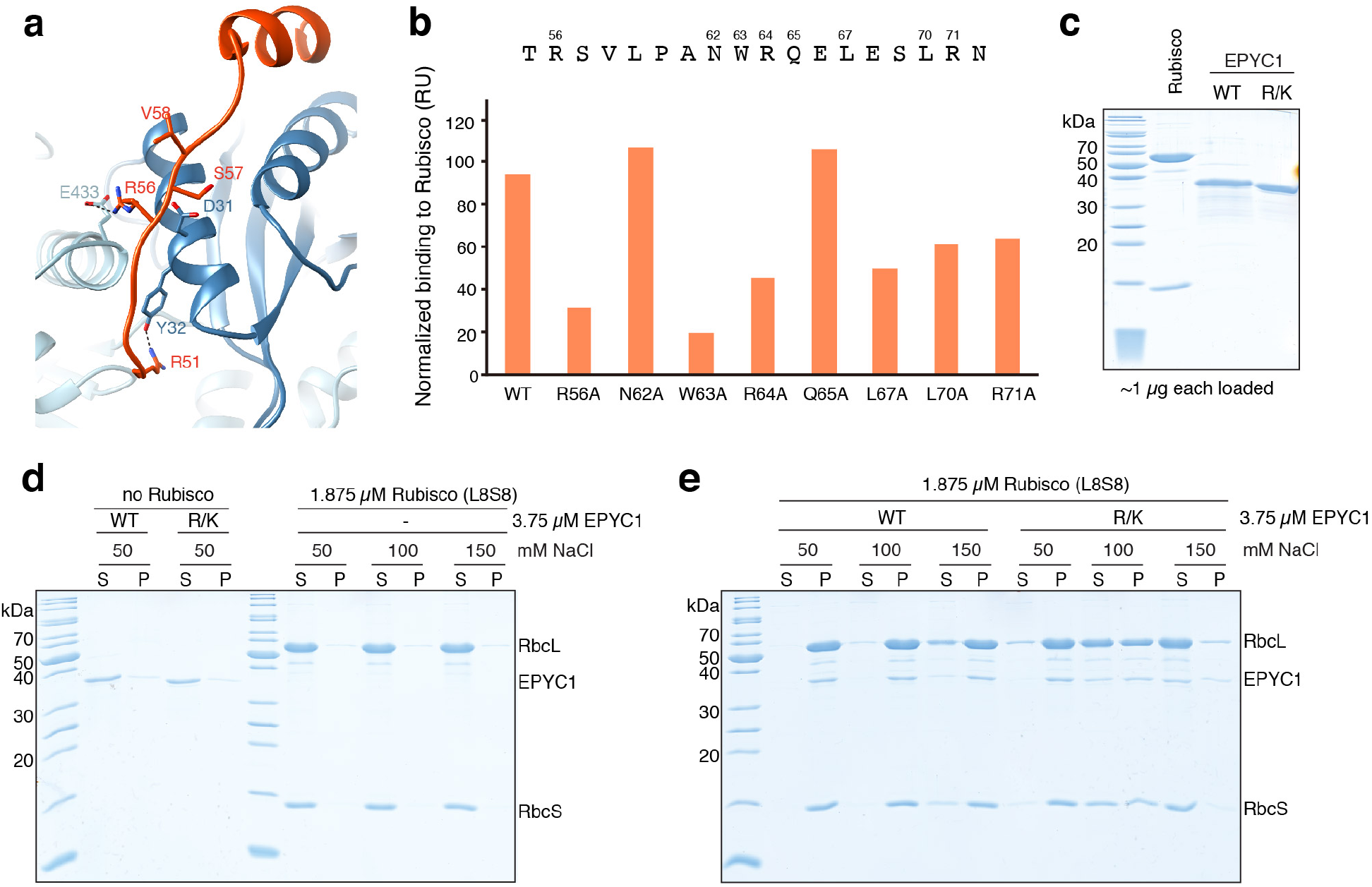
Interface residues on EPYC1 identified by Cryo-EM are important for binding and phase separation of EPYC1 and Rubisco. **a**, Our structure suggests that some residues in addition to the ones shown in Fig. 4 may also contribute to the interaction between EPYC1 and Rubisco. R56 of the EPYC1 peptide may interact with both D31 of the Rubisco small subunit and E433 of the Rubisco large subunit. R51 of the EPYC1 peptide may form a salt bridge with Y32 of the Rubisco small subunit. Residues S57 and V58 of the EPYC1 peptide are close to D31 in the structure, which may explain why replacing either of these residues with a negatively charged residue disrupts binding (Fig. 4a). **b**, The wild-type (WT) EPYC1 peptide or EPYC1 peptides with the indicated point mutations were synthesized, and their Rubisco-binding signal was measured by surface plasmon resonance. **c**, SDS-PAGE analysis of purified proteins used for *In vitro* phase separation experiments. WT = wild-type EPYC1; R/K = EPYC1^R64A/K127A/K187A/K248A/R314^. **d-e**, A droplet sedimentation assay was used as a readout of phase separation complementary to the microscopy analyses shown in Fig. 4b. Proteins at indicated concentrations were mixed and incubated for 10 minutes, then condensates were pelleted by centrifugation. Supernatant (S) and pellet (P) fractions were run on a denaturing gel. The negative controls with no Rubisco or with no EPYC1 are shown in (d), and the wild-type Rubisco with wild-type EPYC1 or mutant EPYC1 are shown in (e).

**Extended Data Fig. 6.**
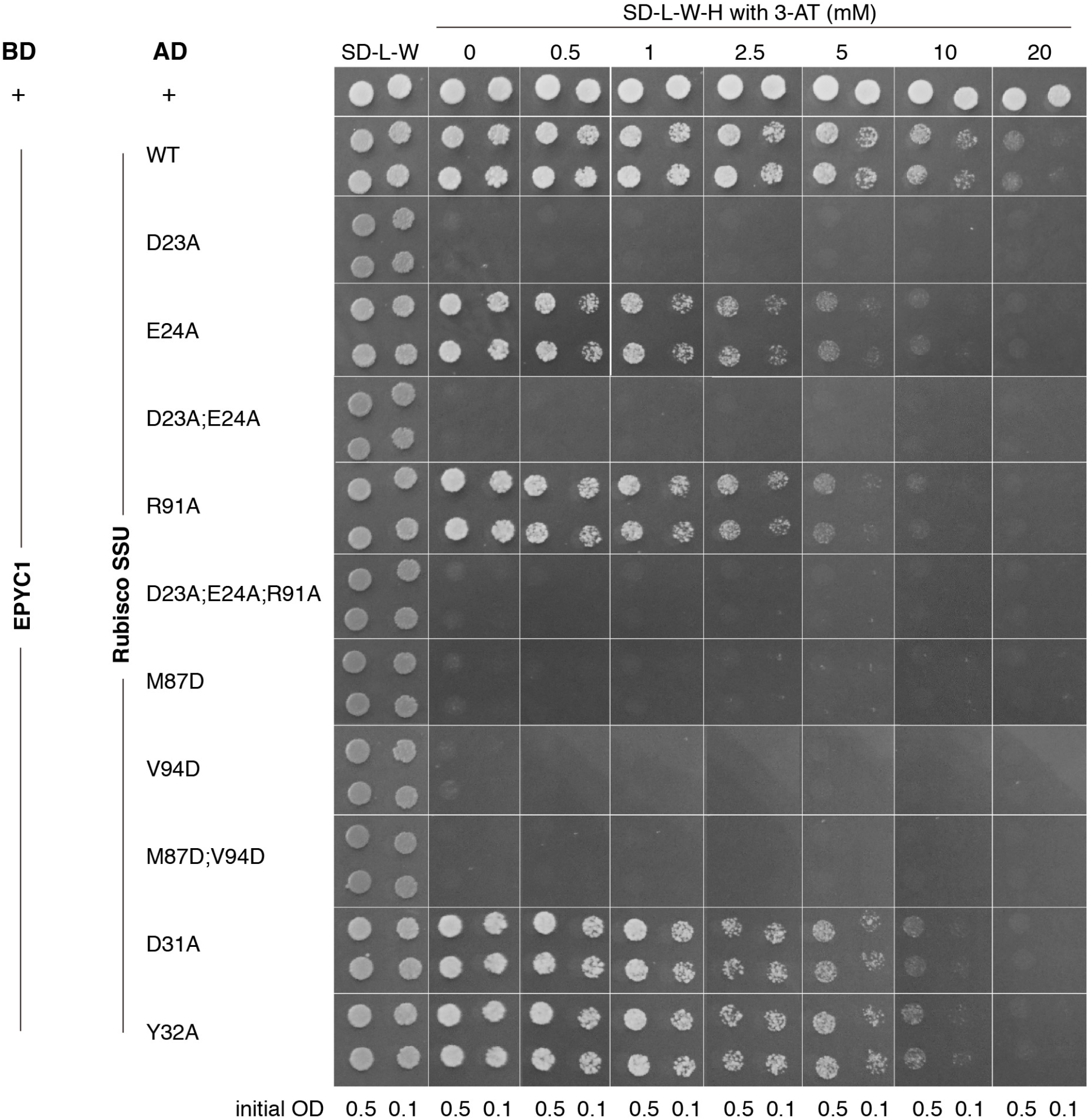
Yeast Two-Hybrid assays of interactions between EPYC1 and wild-type or mutated Rubisco small subunit. Colonies are shown after 3 days’ growth on plates. A subset of the data shown in this figure are shown in Fig. 5a.

**Extended Data Fig. 7.**
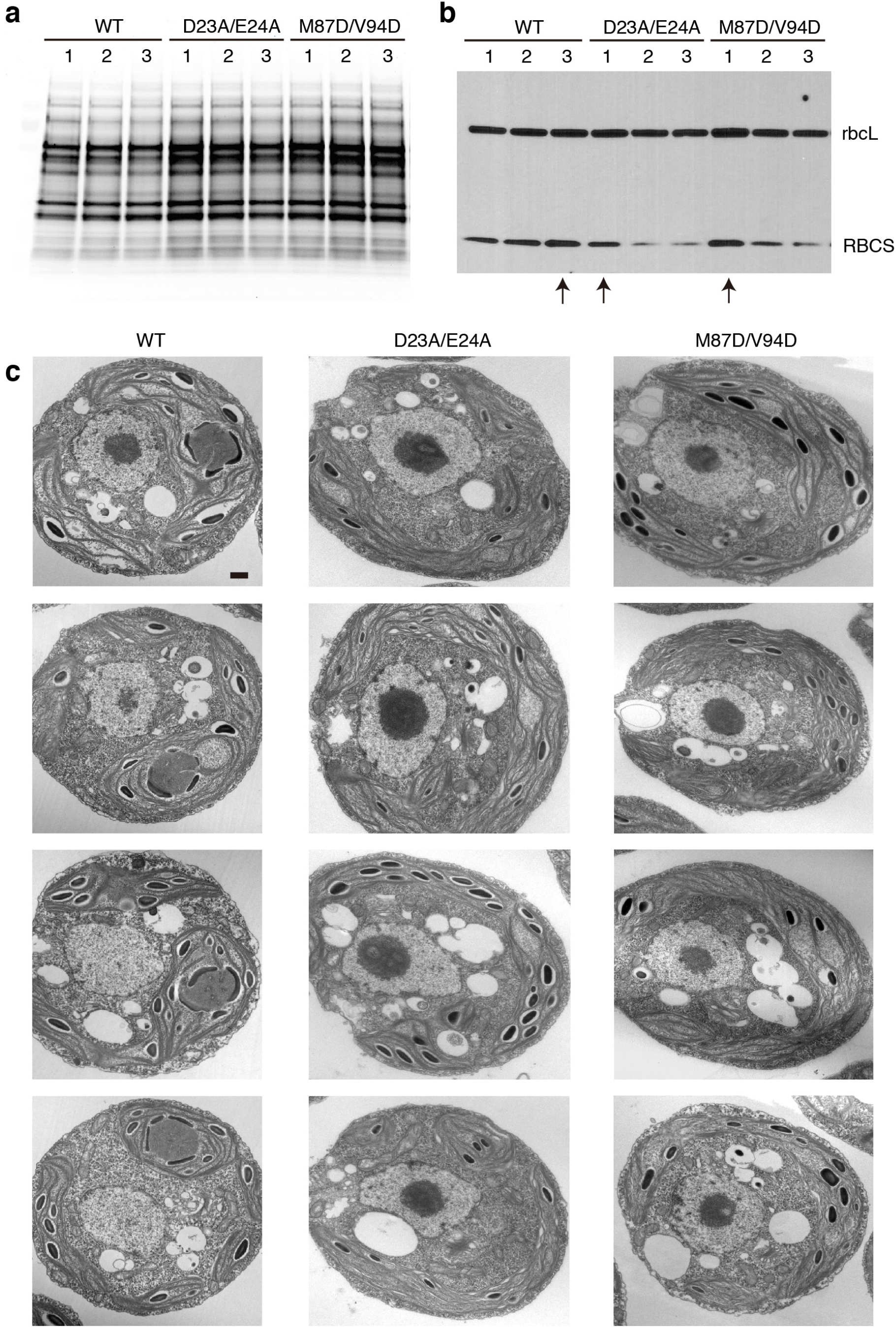
Selection of the Rubisco small subunit mutant strains for phenotype analysis. **a**, The Rubisco small subunit-less mutant T60 (*Δrbcs*) was transformed with DNA encoding wild-type and mutant Rubisco small subunits (RBCS) to produce candidate transformants with the genotypes *Δrbcs;RBCS^WT^, Δrbcs;RBCS^D23A/E24A^*, and *Δrbcs;RBCS^M87D/V94D^*. Total protein extracts for three strains from each transformation were separated on a polyacrylamide gel. **b**, The gel shown in A was probed by Western Blot using a polyclonal antibody mixture that detects both large and small Rubisco subunits. The candidate transformants with highest RBCS expression level from each genotype are indicated by an arrow below the lanes and were used for the subsequent phenotypic analyses shown in Fig. 5 and panel c. **c**, Additional representative TEM images of whole cells of the strains expressing wild-type, D23A/E24A, and M87D/V94D Rubisco small subunit. Scale bar = 500 nm.

**Extended Data Table 1.** Cryo-EM data collection and refinement.

|  | #1 Apo Rubisco<br>(EMDB-xxxx)<br>(PDB xxxx) | 2# EPYC1<sub>49-72</sub> peptide-bound Rubisco<br>(EMDB-xxxx)<br>(PDB xxxx) |
| --- | --- | --- |
| Voltage (kV) | 300 | 300 |
| Magnification | 22,500 | 22,500 |
| Defocus range (μm) | -1.5 to -3.0 | -1.5 to -3.0 |
| Pixel size (Å) | 1.31 | 1.31 |
| Exposure time (s) | 10 | 10 |
| No. movie frames | 50 | 50 |
| Electron dose (e-/Å<sup>2</sup>) | 58 | 58 |
| No. micrographs | 2,500 | 2,500 |
| No. initial particles | 677,071 | 1,809,869 |
| No. final particle | 491,395 | 945,755 |
| Symmetry | D4 | D4 |
| Resolution (Å) | 2.68 | 2.62 |
| Map sharpening *B* factor (Å<sup>2</sup>) | -147.233 | -154.161 |

**Extended Data Table 2.**
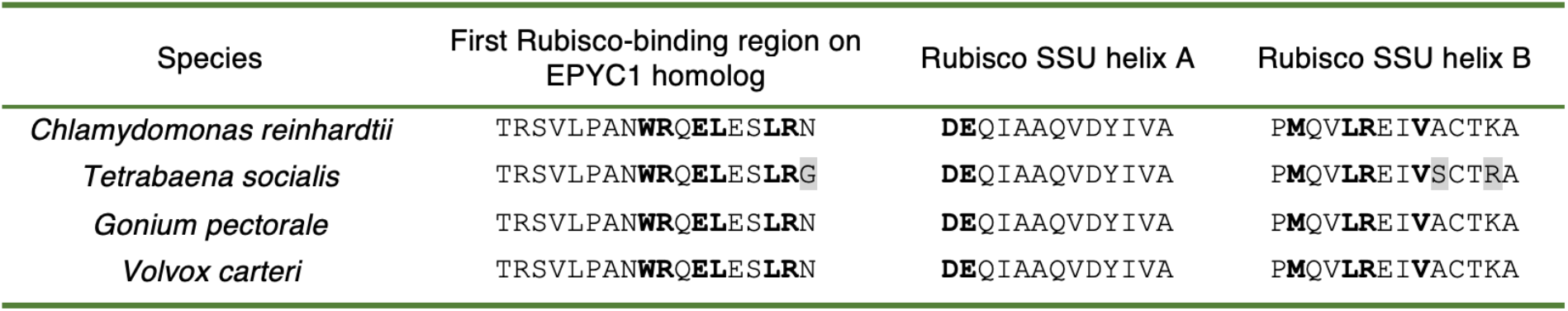
The amino acid residues that form the Rubisco-binding regions on EPYC1 homologs, and the EPYC1 binding site on the surface of Rubisco, appear to be conserved across the order Volvocales. Residues with roles in the binding interface are bolded. Residues that are different from the *Chlamydomonas reinhardtii* sequence are highlighted in grey.

| Species | First Rubisco-binding region on EPYC1 homolog | Rubisco SSU helix A | Rubisco SSU helix B |
| --- | --- | --- | --- |
| *Chlamydomonas reinhardtii* | TRSVLPAN**WRQ**E**LES**L**R**N | **DE**QIAAQVDYIVA | **PMQV**L**REI**V**ACTKA** |
| *Tetrabaena socialis* | TRSVLPAN**WRQ**E**LES**L**R**G | **DE**QIAAQVDYIVA | **PMQV**L**REI**V**SCTRA** |
| *Gonium pectorale* | TRSVLPAN**WRQ**E**LES**L**R**N | **DE**QIAAQVDYIVA | **PMQV**L**REI**V**ACTKA** |
| *Volvox carteri* | TRSVLPAN**WRQ**E**LES**L**R**N | **DE**QIAAQVDYIVA | **PMQV**L**REI**V**ACTKA** |

